# A combination of major histocompatibility complex (MHC) I overexpression and type I interferon induce mitochondrial dysfunction in human skeletal myoblasts

**DOI:** 10.1101/2024.04.10.588847

**Authors:** Anastasia Thoma, Holly L Bond, Tania Akter-Miah, Nasser Al-Shanti, Hans Degens, Vanja Pekovic-Vaughan, Adam P Lightfoot

## Abstract

The overexpression of major histocompatibility complex (MHC) I on the surface of muscle fibres is a characteristic hallmark of the idiopathic inflammatory myopathies (IIMs), collectively termed myositis. Alongside MHC-I overexpression, sub-types of myositis, display a distinct type I interferon (IFN) signature. This study examined the combinational effects of elevated MHC-I and type I IFNs (IFNα/β) on mitochondrial function, as mitochondrial dysfunction is often seen in IIMs. Human skeletal muscle myoblasts were transfected with an MHC-I isoform using the mammalian HLA-A2/K^b^ vector. Mitochondrial respiration, mitochondrial membrane potential, and reactive oxygen/nitrogen species generation were assessed with or without IFNα and IFNβ. We show that MHC-I overexpression in human skeletal muscle myoblasts led to decreased basal glycolysis and mitochondrial respiration, cellular spare respiratory capacity, ATP-linked respiration, and an increased proton leak, which were all exaggerated by type I IFNs. Mitochondrial membrane depolarisation was induced by MHC-I overexpression both in absence and presence of type I IFNs. Human myoblasts overexpressing MHC-I showed elevated nitric oxide generation that was abolished when combined with IFN. MHC-I on its own did not result in an increased ROS production, but IFN on their own, or combined with MHC-I overexpression did induce elevated ROS generation. We present new evidence that MHC-I overexpression and type I IFNs aggravate the effects each has on mitochondrial function in human skeletal muscle cells, providing novel insights into their mechanisms of action and suggesting important implications in the further study of myositis pathogenesis.

## INTRODUCTION

The idiopathic inflammatory myopathies (IIMs) are a group of rare acquired inflammatory muscle diseases, collectively known as myositis. Patients with myositis share some common clinical features, including muscle weakness, increased circulating muscle enzymes (e.g., creatine kinase), inflammatory infiltration of CD4^+^/CD8^+^ T-cells and B-cells within the muscle, expression of muscle-specific and -associated autoantibodies and interferon (IFN) upregulation in muscle fibres, and overexpression of major histocompatibility complex class I (MHC-I) on the surface of muscle fibres [1].

A recent study demonstrated that IFN signature facilitates stratification of IIMs, where for instance elevated expression of both type I IFNα and IFNβ is unique to dermatomyositis [2]. The significance of type I IFN is illustrated by the fact that IFN-I treatment of human peripheral blood mononuclear cells and skeletal muscle cells induced a gene expression profile that was largely identical to that observed in patients with dermatomyositis [3]. IFN-inducible genes were also present in blood samples of patients with dermatomyositis and polymyositis and correlated with disease type. Specifically, dermatomyositis samples presented higher levels of type I IFN-inducible genes compared to polymyositis, while none were found in inclusion body myositis [4].

The distinct type I IFN signature in dermatomyositis as well as the overexpression of MHC-I have turned the attention to their role in downstream pathology-associated mechanisms. Sustained up-regulation of MHC-I in muscle has displayed a positive correlation with endoplasmic reticulum stress activation [5, 30], which has been evident in patients with myositis [6]. Perifascicular atrophy is a histological hallmark of dermatomyositis, and studies have shown that mitochondrial abnormalities and MHC I overexpression, are predominantly localised to these fibres undergoing atrophy [7].

A recent study highlighted the involvement of mitochondrial damage, by the mitochondria-localised pro-apoptotic hara-kiri protein, in myofiber death and muscle weakness in individuals with dermatomyositis and polymyositis [8]. A further study showed decreased expression of genes associated with mitochondrial biogenesis and those encoding proteins of the electron transport chain complexes in muscle fibres from a dermatomyositis patient and in a murine experimental autoimmune model of myositis [9]. This study also demonstrated reduced mitochondrial respiration and increased ROS generation in each model. Furthermore, they provided evidence that the type I IFN score correlated with mitochondrial respiratory deficiency in dermatomyositis muscle, while IFN-β treatment of human myotubes decreased mitochondrial respiration in a ROS-dependent manner [9].

Both type I IFN signature and MHC-I have been suggested to play an important role in dermatomyositis and polymyositis. Despite the evidence of mitochondrial abnormalities in myositis, to our knowledge, the role of sustained MHC-I upregulation on mitochondrial function [31] remains to be revealed. Likewise, little is known about the distinct effects of IFNα and IFNβ on mitochondrial function. Therefore, we have generated an *in vitro* MHC-I human skeletal muscle model that overexpresses human leukocyte antigen (HLA) class I (the so-called MHC-I system in humans) [10]. Using this model, the present study brings insights not only into the role of MHC-I overexpression on mitochondrial function in human muscle cells, but also presents the distinct effects of type I IFNα/β in the presence or absence of MHC-I overexpression. Our data reveals for the first time that MHC-I overexpression in human muscle cells acts synergistically with type I IFNs resulting in mitochondrial respiratory defects and that each on their own present distinct effects on reactive oxygen and nitrogen species production.

## MATERIALS AND METHODS

### Cell culture

An immortalised human skeletal muscle cell line was a gift from the Centre for Research in Myology in Paris (France) [11]. Cells were grown under standard cell culture conditions (37°C, 5% CO_2_) in growth media (GM) containing: Dulbecco’s Modified Eagles Medium, high-glucose (4.5 g/L DMEM, Lonza, UK) and Medium-199 with Earle’s BSS (1:5, v/v) (Sigma-Aldrich, UK), 20% (v/v) heat inactivated foetal bovine serum (Gibco, UK), 1% (v/v) penicillin/streptomycin, 1% (v/v) L-glutamine (Lonza, UK), 10 μg/mL gentamicin, 25 ng/mL fetuin from foetal bovine serum, 0.2 μg/mL dexamethasone, 5 μg/mL recombinant human insulin (Sigma-Aldrich, UK), 0.5 ng/mL recombinant human basic fibroblast growth factor, 5 ng/mL recombinant human epidermal growth factor, and 2.5 ng/mL recombinant human hepatocyte growth factor (Gibco, UK).

### Plasmid preparation

Psv2-neo plasmid containing HLA-A2/K^b^ was a gift from Linda Sherman (Addgene #14906, USA) [12]. Psv2-neo empty vector (EV) plasmid in *Escherichia coli* was purchased from ATCC (#37149, UK). The plasmid DNA (pDNA) was amplified by creating single colonies of *Escherichia coli* in agar plates and incubated in Lennox L Broth (LB Broth; Invitrogen, UK). The pDNA was stored as glycerol stocks and purified using QIAprep spin miniprep kit (Qiagen, UK). The concentration was determined before use and only plasmid DNA with 260/280 > 1.80 and 260/230 > 2.0 was used.

### Transfection and treatments

Human skeletal muscle myoblasts were seeded 24 hours before transfection to achieve 70–90% confluence. At the time of transfection, the GM was replaced with fresh GM. Cells were incubated with the transIT-X2 (Mirus, USA): plasmid (EV or HLA-A2/K^b^; 1 µg/µL) complex at 2:1 ratio prepared in Opti-MEM (Gibco, UK) for 18 hours in the presence or absence of 100 ng/mL type I IFNα (PBL Assay Science, USA) or IFNβ (R&D Systems, USA).

### Assessment of transfection efficiency by immunostaining

Transfection efficiency was determined by immunostaining. Human skeletal muscle myoblasts were fixed in 4% paraformaldehyde (Alfa Aesar, USA) (15 minutes, room temperature) and permeabilised using 0.5% Triton X-100 (Sigma, UK) (15 minutes, room temperature). Cells washed in Dulbecco’s Phosphate-Buffered Saline (DPBS, Lonza, UK) were blocked using 3% goat serum (Vector Laboratories, USA) in 0.05% Tween-20 (Fisher Scientific, UK) in PBS for 1 hour at room temperature. Washed cells were incubated with anti-HLA class I antibody (1/1000, Abcam #ab23755, UK) at 4^D^C overnight and stained with goat anti-mouse IgG (H+L) Alexa Fluor 488 (1/800, Invitrogen, UK) and 4′,6′-diamidino-2-phenylindole dihydrochloride (DAPI, 1/3000, Sigma-Aldrich, UK) for 1 hour at room temperature protected from ambient light. Cells maintained in DPBS were imaged using a LEICA DMI6000 B inverted microscope (CTR6000 laser, Leica Microsystems).

### Assessment of cellular respiration using Seahorse Extracellular Flux Analysis

Mitochondrial function and glycolytic activity were measured with an XFp Extracellular Flux Analyser (Agilent Technologies, UK). Human skeletal muscle cells were seeded in an 8-well XFp plate at a density of 7×10^3^ cells/well in GM. After overnight incubation, cells were transfected with the HLA-A2/K^b^ overexpression vector or EV in the presence or absence of type I IFNα or IFNβ, as described above. Real-time oxygen consumption rate (OCR) and extracellular acidification rate (ECAR), as a measure of mitochondrial respiration and basal glycolysis, respectively [13], were assessed using the Seahorse XFp Mito Stress Test (Agilent Technologies, UK). GM was replaced with Seahorse assay medium containing unbuffered DMEM (pH 7.4), 1 mM pyruvate (Agilent Technologies, UK), 2 mM L-glutamine, and 10 mM glucose (Agilent Technologies, UK), and myoblasts were incubated at 37°C in a non-CO_2_ incubator 1 hour prior the experiment. The sequential addition of oligomycin (1 μM), FCCP (2 μM), and rotenone/antimycin (0.5 μM) enabled the analysis of the cell bioenergetic phenotype parameters. Oligomycin, an ATP synthase inhibitor, was injected to enable measurement of ATP-linked respiration. This also allowed assessment of proton leak, as the basal respiration that is not used for ATP production. The uncoupling agent FCCP was added to disrupt OCR and ECAR, normalised to total protein concentration using the Pierce™ BCA Protein Assay (Thermo scientific, Loughborough, UK), calculated by Seahorse XFp Wave software version 2.2.0 (Agilent Technologies, UK).

### Assessment of mitochondrial membrane potential and mass

Human skeletal muscle myoblasts were seeded at 8×10^3^ cells/well in a black-sided, clear-bottom microplate (96 wells), cultured in GM, and transfected and treated as described above. JC-1 fluorophore (5,5′,6,6′-tetrachloro-1,1′,3,3′-tetraethylbenzimi-dazolylcarbocyanine iodide) (Abcam, Cambridge, UK), MitoTracker Red CMXRos and TMRM (Tetramethylrhodamine, methyl ester) (Molecular Probes, Invitrogen, Paisley, UK) were used to assess mitochondrial membrane potential (ΔΨm). Human skeletal muscle myoblasts were incubated with JC-1 (5 µM, 30 minutes, 37°C) in the presence of transfection solution (HLA-A2/K^b^ or EV) with or without type I IFNs, in the dark. Following incubation with JC-1, cells were washed with DPBS and maintained in GM containing type I IFN treatments, and fluorescence intensities from JC-1 aggregate and monomer forms were measured at excitation 530/25 and 485/20 nm, respectively, and emission 590/35 nm. Changes in TMRM fluorescence intensity were normalised to mitochondrial mass by washing myoblasts transfected with HLA-A2/K^b^ or EV with or without type I IFNs with warm DBPS and incubating them with TMRM/MitoTracker Green FM (Invitrogen, UK) (10 nM and 100 nM, respectively, 30 minutes, 37°C). Fluorescence intensity was read in the presence of staining solution at 530/25 nm excitation and 590/35 nm emission for TMRM and 485/20 nm excitation and 528/20 nm emission for MitoTracker Green FM. To assess MitoTracker Red CMXRos fluorescence intensity, myoblasts were washed with warm DPBS and stained in 5 µM MitoTracker Red CMXRos solution (30 minutes, 37 °C) and fluorescence was measured in cells maintained in phenol red-free DMEM at excitation 590/20 and emission 645/40 nm. Endpoint fluorescence for all probes was measured using a SynergyTM multi-detection microplate reader (BioTek Instruments, UK). All measurements were corrected for background fluorescence and normalised to total protein content.

### Measurement of reactive oxygen/nitrogen species

To quantify mitochondrial superoxide and hydroxyl radical generation, human myoblasts transfected with HLA-A2/K^b^ or EV +/- type I IFNs were seeded in a black-sided, clear-bottom microplates (96 wells), washed with DBPS and incubated with either a MitoSOX™ Red mitochondrial superoxide indicator (5 µM, 30 minutes, 37°C) (Invitrogen, UK) or OH580 probe, a mitochondrial hydroxyl radical indicator (1 hour, 37°C) (Abcam, UK) in phenol red-free DMEM in the dark. Following incubation, cells were washed with DPBS, maintained in assay medium, and endpoint fluorescence was measured with the following excitation and emission wavelengths: MitoSOX™ Red, 360/40 and 590/35 nm; and OH580 probe, 530/25 and 590/35 nm, respectively. To assess intracellular nitric oxide generation, DAF-FM-DA (10 µM) (Abcam, UK) was added to the cells in the presence of transfection solution +/- type I IFNs. Following 30 minutes incubation at 37°C, cells were washed with DPBS, maintained in GM +/- type I IFNs, and endpoint fluorescence was measured at 485/20 nm excitation and 528/20 nm emission. Amplex® Red Hydrogen Peroxide/Peroxidase assay (Invitrogen, UK) was used to measure hydrogen peroxide (H_2_O_2_) released from cells according to the manufacturer’s instructions. Briefly, cell culture media and standard curve samples of known H_2_O_2_ concentrations were incubated with the Amplex® Red reagent (100 µM)/Horseradish peroxidase (0.2 U/mL) solution) at room temperature for 30 minutes and endpoint fluorescence was measured at 530/25 nm excitation and 590/35 nm emission. Endpoint fluorescence for all probes was measured using a SynergyTM multi-detection microplate reader (BioTek Instruments, UK). All measurements were corrected for background fluorescence and normalised to total protein content.

### Statistical analyses

Statistical analyses were performed with GraphPad Prism version 8. Data were assessed for normality of distribution using Shapiro-Wilk test. Normally distributed data were analysed using one-way ANOVA followed, where appropriate, by Tukey’s post hoc test, and non-normally distributed data using Kruskal-Wallis test with Dunn’s post hoc test. A *p*-value ≤ 0.05 was considered statistically significant.

## RESULTS

### IFN**α** and IFN**β** present a differential impact on mitochondrial respiration

We initially examined the effects of the type of IFNs on the mitochondrial function of human skeletal muscle cells using Seahorse Extracellular Flux Analyser. Eighteen hours exposure to 100 ng/mL or 200 ng/mL IFNα treatment did not induce any significant changes in mitochondrial function parameters, including spare respiratory capacity, basal, maximal, ATP-linked, and leak respiration, nor did it affect non-mitochondrial respiration (**Figure 1A-F**). Mitochondrial bioenergetics remained unaffected following 24 hours treatment with 100 ng/mL IFNα, while higher IFNα concentration (200 ng/mL) significantly reduced oxygen consumed for basal and ATP-linked respiration only, indicating the initiation of mild effects on mitochondrial function by a longer exposure of human muscle cells to IFNα (**Figure 1G-L**).

**Figure 1.**
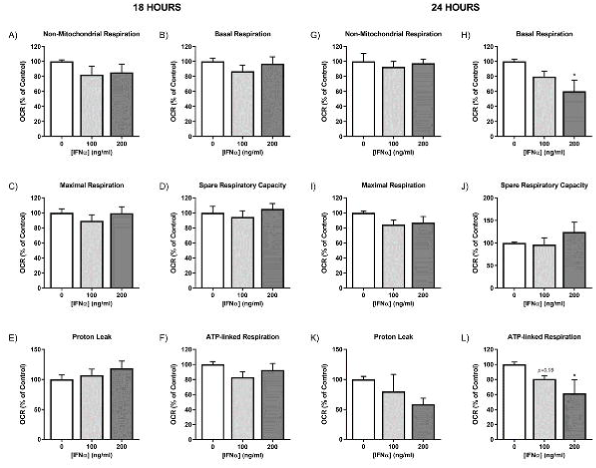
Mitochondrial function of human myoblasts treated with interferon-α (IFNα). Non-mitochondrial respiration, basal respiration, maximal respiration, spare respiratory capacity, proton leak, and ATP-linked respiration, normalised to protein content following 18 hours (**A–F**) and 24 hours (**G–L**) incubation with IFNα at indicated doses (*n* = 6). Data represent mean ± S.E.M., \**p* ≤ 0.05 compared to control.

In contrast to IFNα treatment, treatment of human myoblasts with 100 ng/mL or 200 ng/mL of IFNβ for 18 hours induced a significant reduction in basal, maximal, and ATP-linked respiration, while defects in non-mitochondrial respiration were induced by 200 ng/mL IFNβ only (**Figure 2B-F**). It should be noted that 18 hours exposure to IFNβ at either concentration did not induce changes in spare respiratory capacity or leak respiration (**Figure 2D and 2E**). Interestingly, lower proton leak was seen following 24 hours treatment with IFNβ (**Figure 2K)**, with no discernable changes being evident in other mitochondrial function parameters (**Figure 2G-L**).

**Figure 2.**
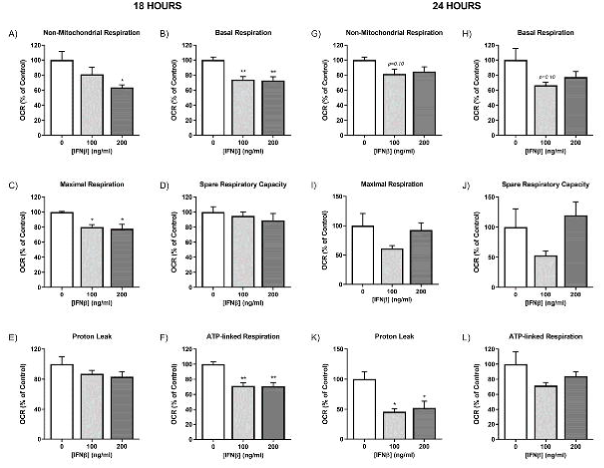
Mitochondrial function of human myoblasts treated with interferon-β (IFNβ). Non-mitochondrial respiration, basal respiration, maximal respiration, spare respiratory capacity, proton leak, and ATP-linked respiration, normalised to protein content following 18 hours (**A–F**) and 24 hours (**G–L**) incubation with IFNβ at indicated doses (*n* = 6). Data represent mean ± S.E.M., \**p* ≤ 0 .05, \*\**p* < 0.01 compared to control.

### MHC-I levels following transfection with HLA-A2/K^b^ vector in the presence or absence of type I IFNs

We next transfected human skeletal cells with either empty vector EV or HLA-A2/K^b^ vectors and examined their expression using fluorescence microscopy. Transfection of human skeletal myoblasts with the HLA-A2/K^b^ vector resulted in significantly upregulated MHC-I expression compared to empty vector as shown by changes in fluorescence intensity levels of HLA (green channel fluorescence) normalised to nuclei number (blue channel fluorescence) (**Figure 3**). Interestingly, IFNα exposure of MHC-I overexpressing human myoblasts induced significantly higher HLA fluorescence intensity levels (by 78.2%), compared to HLA-transfected myoblasts alone. In contrast, IFNβ exposure of MHC-I overexpressing human myoblasts increased HLA fluorescence intensity only by 20.5%, compared to MHC-I expression alone (**Figure 3A and 3B**).

**Figure 3.**
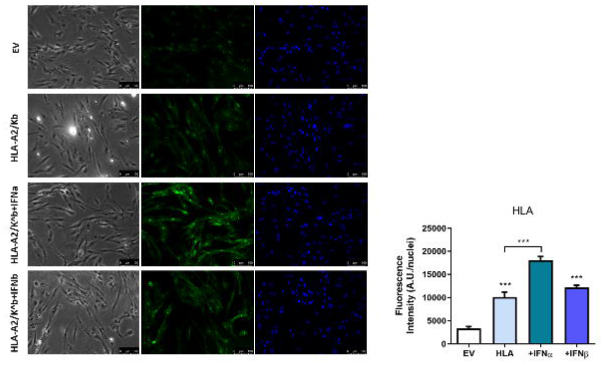
*In-vitro* overexpression of major histocompatibility complex-I (MHC-I). (**A)** Representative phase contrast and single-channel fluorescent images of human skeletal muscle myoblasts transfected with HLA-A2/K^b^ or empty vector (EV) in presence or absence of type I IFNs stained with anti-HLA-I (green) and DAPI (blue). Images captured at 20× magnification. Scale bar = 100 μm. (**B**) Quantification of HLA Class I fluorescence intensity level normalised to nuclei number. Data represent mean ± S.E.M., \**p* ≤ 0.05, \*\*\**p* < 0.001 compared to empty vector or to HLA I-overexpressing cells.

### MHC-I overexpression-induced changes in cellular metabolism are exacerbated by type I IFNs

Overexpression of MHC-I in human skeletal muscle myoblasts induced a reduction in both mitochondrial respiration (OCR) and basal glycolysis (ECAR) as assessed by Seahorse Extracellular Flux Analyser, which was exacerbated in the presence of 100 ng/ml type I IFNs (IFNα or IFNβ) following 18 hours exposure (**Figure 4A-D**). MHC-I overexpressing myoblasts in the presence of IFNβ also showed a significantly reduced non-mitochondrial respiration (**Figure 4E**). Mitochondrial OCR was suppressed at both basal and maximal capacities by MHC-I overexpression, which was exacerbated in the presence of IFNα or IFNβ, with IFNβ showing stronger effects (**Figure 4F and 4G**). Decreased mitochondrial respiration was accompanied by significantly decreased ATP-linked respiration in MHC-I overexpressing human myoblasts, with IFNβ inducing the largest decrease, followed by IFNα, when compared to EV (**Figure 4I**). IFNβ treatment of MHC-I overexpressing myoblasts also significantly decreased spare respiratory capacity (**Figure 4H**). Moreover, proton leak respiration was decreased by MHC-I overexpression, with combinatorial IFNα or IFNβ treatments leading to a higher rate of change, when compared to EV (**Figure 4J**).

**Figure 4.**
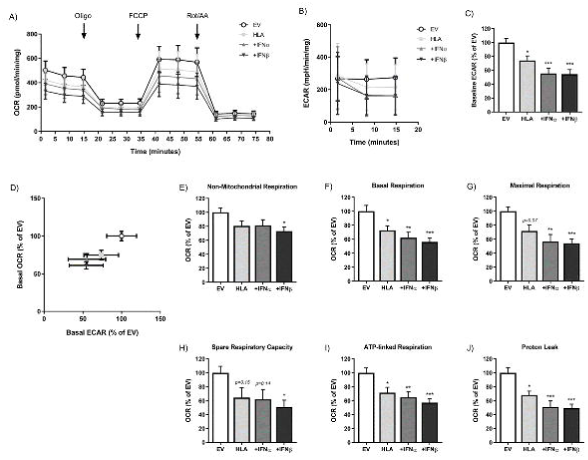
Mitochondrial function of HLA-I-transfected human myoblasts treated with or without type I interferons (IFNs). (**A**) Real-time measurements of oxygen consumption rate (OCR) following the sequential injection of oligomycin, FCCP, and a mixture of rotenone/antimycin A. (**B**) Real-time measurements of extracellular acidification rate (ECAR) and (**C**) baseline ECAR values. (**D**) Bioenergetic profile expressed as OCR versus ECAR measured under basal condition. (**E–J**) Mitochondrial function parameters; non-mitochondrial respiration, basal respiration, maximal respiration, spare respiratory capacity, ATP-linked respiration, and proton leak, normalised to protein content (*n* = 8 per group). Data represent mean ± S.E.M., \**p* ≤ 0.05, \*\**p* < 0.01, \*\*\**p* < 0.001 compared to empty vector.

### MHC-I overexpression in the presence of type I IFNs induces changes in mitochondrial mass and membrane potential

Different fluorophores were used as probes to assess changes in mitochondrial membrane potential in response to MHC-I overexpression of human myoblasts in the presence or absence of type I IFNs. We first determined the mitochondrial membrane potential using the ratiometric dye JC-1, which reflects the ratio of JC1-aggregates formed in active (negatively charged) mitochondria *vs.* JC1-monomers present in depolarised mitochondria [14]. The JC-1 ratio was significantly decreased in MHC-I overexpressing human muscle cells, with type I IFNs inducing a higher rate of decrease by approximately 35% compared to MHC-I overexpressing cells alone (**Figure 5A**). We also utilised another mitochondria-based dye, TMRM, which accumulates solely in active mitochondria with intact membrane potential. MHC-I overexpression in combination with IFNα, but not IFNβ, induced a significant increase in TMRM fluorescence (**Figure 5C**). MHC-I overexpression in combination with IFN, but not on its own, induced a significant increase in mitochondrial mass as assessed by MitoTracker Green FM fluorescence intensity (**Figure 5D**). However, when the TMRM fluorescence signals were normalised to mitochondrial mass using MitoTracker Green FM, the TMRM fluorescence intensity was reduced, rather than increased in MHC-I overexpressing cells in the presence of IFNβ treatment (**Figure 5E**). The MitoTracker Red CMXROS probe, which accumulates in active mitochondria, showed no significant change in fluorescence intensity in MHC-I myoblasts both in the absence and presence of type I IFNs (**Figure 5B**).

**Figure 5.**
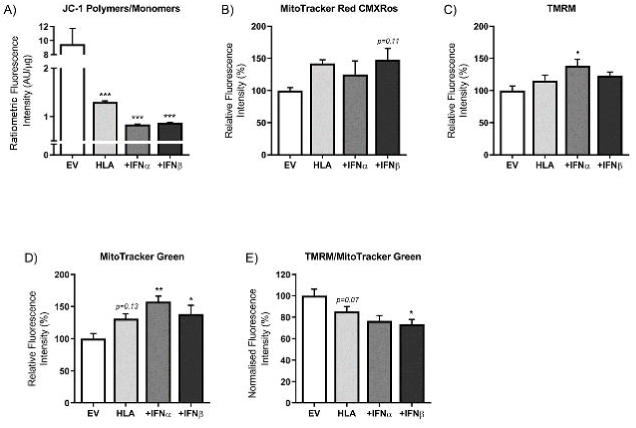
Mitochondrial mass and membrane potential of HLA-I-transfected human myoblasts treated with or without type I interferons (IFNs). (**A**) Mitochondrial membrane potential expressed as JC-1 aggregates (red fluorescence) to JC-1 monomers (green fluorescence) ratio normalised to protein content (*n* = 6). (**B–D**) Fluorescence intensity changes of MitoTracker Red CMXRos and TMRM indicative of changes in mitochondrial membrane potential of active mitochondria only, as well as of MitoTracker Green, representing changes in mitochondrial mass. (**E**) Fluorescence intensity of TMRM normalised to MitoTracker Green. Data represent mean ± S.E.M., \**p* ≤ 0.05, \*\**p* < 0.01, \*\*\**p* < 0.001 compared to empty vector.

### MHC-I overexpression in human myoblasts leads to increased nitric oxide generation, while mitochondrial superoxide is elevated in the presence of combined treatment with IFN**β**

We then examined RONS production in MHC-I overexpressing human myoblasts in the absence or presence of type I IFNs. Type I IFNs alone led to a significant increase in mitochondrial superoxide production as assessed by MitoSOX Red compared to control myoblasts (**Figure 6A**), while MHC-I overexpressing cells did not show an elevated ROS production (**Figure 6B**). An increase in MitoSOX Red fluorescence intensity in MHC-I-overexpressing myoblasts was only observed in the presence of type I IFNβ (**Figure 6B**), while no significant change was observed in the case of IFNα treatment (**Figure 6B**).

**Figure 6.**
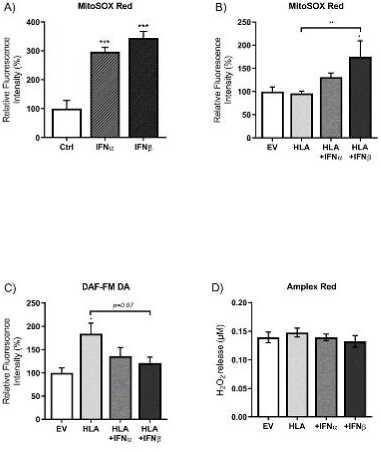
RONS generation in HLA-I-transfected human myoblasts treated with or without type I IFNs. (**A–B**) Fluorescence intensity levels of MitoSOX Red, an indicator of mitochondrial superoxide generation, induced by type I IFNs or MHC-I overexpressing cells with or without type I IFNs (*n* = 3**–**8), and (**C**) DAF-FM DA, showing cellular nitric oxide generation (*n* = 4),. (**D**) Release of hydrogen peroxide from cells as measured by Amplex Red assay. All data was normalised to protein content. Data represent mean ± S.E.M., \**p* ≤ 0.05, \*\**p* < 0.01, \*\*\**p* < 0.001 compared to empty vector or HLA I-overexpressing cells.

Overexpression of MHC-I alone promoted an increased intracellular NO generation in human myoblasts compared to the EV-expressing myoblasts, indicated by the increased DAF-FM DA fluorescence intensity (**Figure 6C**). The NO levels were also substantially higher in MHC-I overexpression alone compared to MHC-I overexpression + IFNβ treatment (**Figure 6C**). Lastly, there was no change in H_2_O_2_ release from human myoblasts as assessed by Amplex Red following MHC-I overexpression in the absence or presence of type I IFNs compared to EV-expressing myoblasts (**Figure 6D**).

## DISCUSSION

Previous research has shown that MHC-I overexpression can induce ER stress, potentially driving skeletal muscle weakness in myositis in the absence of inflammatory infiltrates in muscle fibres [15, 16]. Distinct IFN signatures have been suggested to act as biomarkers of different IIMs, with type I IFNs being predominantly seen in dermatomyositis and to a lesser degree in polymyositis [17]. Given the bi-directional crosstalk between ER and mitochondria, as well as evidence of mitochondrial abnormalities in the muscle of myositis patients [7–9, 18], this study aimed to assess the effects of MHC-I overexpression in human skeletal muscle myoblasts in the presence or absence of type I IFNs on mitochondrial functionality. The major findings of this research are that the negative impact of MHC-I overexpression on mitochondrial function membrane potential and aggravated by the presence of type IFNs, alongside a differential impact on RONS generation.

In this study, we observed that IFNα induced a further overexpression of MHC-I in the HLA-transfected human skeletal muscle model. This finding is consistent with previous studies showing IFNα induced long-lasting MHC-I overexpression in human beta cells [19, 20]. Moreover, a study in human primary skeletal muscle cells found that upregulated levels of HLA-ABC-induced type I IFN pathway activation, which inhibited myoblast differentiation and induced myotube atrophy in the context of diabetes [21]. Here, we were able to reveal that it is IFNα, rather than IFNβ, that induced enhanced HLA-I expression in MHC-I overexpressing cells, highlighting the differential effects of IFNα and IFNβ on MHC class I regulation in human skeletal myoblasts.

Whilst IFNα enhanced the expression of MHC-I, IFNβ induced stronger defects on mitochondrial function compared to IFNα, independent of MHC-I overexpression. More specifically, IFNα induced mild mitochondrial respiratory defects, specifically seen in basal and ATP-linked respiration, that only became evident after 24 hours incubation at a concentration of 200 ng/mL, while such effects were already evident after 18 hours incubation in the presence of 100 ng/mL IFNβ. This is consistent with IFNβ-induced mitochondrial damage reported in previous studies of brown adipose tissue [22] and skeletal muscle [9]. However, the IFNβ-induced mitochondrial dysfunction can be considered moderate, as IFNβ seems to decrease basal, maximal, and ATP-linked respiration, but not reserve capacity, and such effects were returned to control values with longer exposure (24 hours).

Our data show direct negative effects of MHC-I overexpression on mitochondrial respiration, as well as basal glycolysis, in human skeletal muscle cells. Interestingly, these effects were exacerbated in presence of type I IFNs. Mitochondrial impairments were also evident as mitochondrial membrane depolarisation in response to MHC-I overexpression in combination with type I IFNβ. Further, a combinational effect of MHC-I overexpression and IFNβ increased mitochondrial mass, which may be interpreted as an attempt by the cells to compensate for respiratory deficits. Interestingly, Kissig *et al*. (2017) reported in brown adipocytes no IFNα-induced changes increase in mitochondrial biogenesis [22], but IFNβ was not studied. Although we cannot draw a firm conclusion, the observed increase in mitochondrial mass in our study may be the result of a synergistic action of MHC-I overexpression and type I IFN. Further studies are warranted to examine the effects of individual type I IFNs in this matter. Altogether, these findings suggest that there is a synergistic impact of MHC-I overexpression and type I IFNs on mitochondrial biogenesis, function, and membrane polarization in human muscle myoblasts.

Reactive oxygen and nitrogen species have been reported to play an important role in the pathogenesis of several myopathies other than myositis. Hence it is possible that reactive oxygen and nitrogen species are involved in myositis-associated muscle weakness [23]. Indeed, the involvement of nitric oxide in inflammatory myopathies has been reported by Tews and Goebel (1998), who showed inducible and neuronal nitric oxide synthase up-regulation in the muscle fibres of 21 patients with myositis [26]. Another study has also shown elevated inducible nitric oxide synthase and nitric oxide levels, as assessed by nitrated tyrosine residue staining, in the muscle fibres of sporadic inclusion body myositis and to a lower degree in dermatomyositis [27]. However, the role of reactive oxygen and nitrogen species in myositis are poorly understood. The present study showed that both IFNα and IFNβ contribute to mitochondrial ROS generation, as indicated by increases in mitochondrial superoxide levels, whilst MHC-I overexpression alone showed no significant increase in mitochondrial superoxide generation. This observation is consistent with a previous study showing IFNβ-induced reactive oxygen species generation in a dose-dependent manner [9]. In our study, mitochondrial superoxide generation was also highly induced in MHC-I overexpressing myoblasts in the presence of type I IFNs, with IFNβ-treated MHC-I overexpressing cells showing statistically significant increase. It was surprising, however, that IFNα-treated MHC-I overexpressing cells showed no significant increase in mitochondrial superoxide generation, considering pronounced elevations observed by IFNα treatment alone. Surprisingly, IFNβ-treated MHC-I overexpressing cells showed a less robust increase in mitochondrial superoxide compared to IFNβ-treated cells only. One explanation for this unexpected observation is that previous studies have shown that nitric oxide can react with superoxide producing peroxynitrite, which may result in an overall decreased superoxide availability [24, 25] and the increased levels of nitric oxide during MHC-I overexpression may have scavenged the oxygen radicals that have been generated as a result of IFN. Alternatively, these data may suggest MHC I overexpression has a blunting effect on mitochondrial superoxide generation, when used in combination with type I IFNs. Moreover, we observed a decline in DAF-FM DA (marker of NO^-^ generation) fluorescence in the presence of IFNs, compared with MHC I overexpression alone. Thus, overall, these findings highlight a potential differential impact of MHC I overexpression and type I IFNs on ROS generation. Further investigations of ROS generation in our study showed no changes in the release of hydrogen peroxide in MHC-I overexpressing cells in presence or absence of type I IFNs. In further support of these findings, both IFNα and IFNβ showed inhibitory effects on nitric oxide production in endothelial cells [28, 29].

There are limitations to the present study, as our findings are obtained in a human muscle cell model of myositis and need to be verified in human patient cells. Thus, although MHC-I overexpression is a key hallmark of myositis, the magnitude of overexpression in our model may not be representative of the levels of MHC I expressed on the surface of a patient’s muscle cells. Similarly, the levels of IFNs we utilised *in vitro* may not reflect the levels seen in muscles from patients.

The present findings provide novel insights into the differential effects of two major type I IFNs, namely IFNα and IFNβ, on mitochondrial respiration, membrane polarisation and a differential impact on RONS generation. This study also shows the effects of MHC-I overexpression and type I IFNs specifically on mitochondrial function, acting synergistically in inducing respiratory defects and a loss of mitochondrial membrane potential. Moreover, it is of great interest that MHC-I overexpression and type I IFNs exert distinct effects on reactive oxygen and nitrogen species production, with MHC-I overexpression contributing to nitric oxide generation but type I IFNs alleviating this effect, and IFNs promoting mitochondrial superoxide production. Hence, these findings have important implications in myositis pathogenesis and warrant further investigation.

## ACKNOWLEDGMENTS

We would like to thank Dr Joanna Parkes for helpful discussion and comment on the data and manuscript.

## AUTHOR CONTRIBUTIONS

Conception and experimental design: Adam Lightfoot, Anastasia Thoma; Data collection: Anastastia Thoma, Holly Bond, Tania Akter-Miah; Data analysis & interpretation: Adam Lightfoot, Anastasia Thoma, Nasser Al-Shanti, Hans Degens, Vanja Pekovic-Vaughan; Manuscript writing: Adam Lightfoot, Anastasia Thoma, Nasser Al-Shanti, Hans Degens, Vanja Pekovic-Vaughan; Final approval of manuscript: All authors.

